# Therapeutic Inhibition of Metalloproteases by Tetracyclines during Infection by Multi-Drug Resistant *Pseudomonas*

**DOI:** 10.1101/2025.02.14.638345

**Authors:** Mary Carr, Sophia R. Chou, Joshua Sun, Doris L. LaRock, Christopher N. LaRock

## Abstract

**Introduction:** Both host and microbial proteases contribute to the destruction of tissue during *Pseudomonas aeruginosa* pulmonary infections. We previously reported that beneficial therapeutic effects of metalloprotease inhibitors in *Pseudomonas aeruginosa* pulmonary infections are not only from targeting the host matrix metalloproteases, the intended target, but also from cross-inhibition of the secreted bacterial metalloprotease LasB.

**Methods:** We used computation modeling and biochemical assays to investigate whether doxycycline and other structurally-related antibiotics of the tetracycline family inhibit the *Pseudomonas* protease, LasB. Modeling infection *in vitro* with human neutrophils and *in vivo* with a murine model of pulmonary infection, we investigate the effect of on tetracyclines inhibit LasB activity, bacterial survival, and inflammation.

**Results:** Tetracycline, doxycycline, minocycline, and sanocycline all inhibit proteolysis by *Pseudomonas* LasB. At sub-antimicrobial concentrations, this sensitizes the bacterium killing by neutrophils. During pulmonary infection, tetracyclines significantly reduce pathological inflammation in the lung.

**Conclusions:** Our study shows that tetracycline-family antibiotics, known inhibitors of mammalian matrix metalloproteinases, also inhibit LasB and have beneficial anti-inflammatory activity during infections by extensively drug-resistant *P. aeruginosa*. By controlling pathological host inflammation, inhibiting bacterial virulence factors, and directly killing bacteria, tetracyclines can have tripartite therapeutic benefits.

## INTRODUCTION

Antibiotics have been a mainstay in modern medicine, but the emergence and increasing prevalence of multidrug resistance (MDR) has begun to limit their utility. This challenge is evident with *Pseudomonas aeruginosa*, a ubiquitous opportunistic pathogen capable of causing severe soft-tissue and pulmonary infections. Isolates are now increasingly reported that are refractory to all conventional antibiotic therapy, making it a World Health Organization and U.S. Centers for Disease Control priority pathogen for therapeutics development (1).

Emergent strategies against MDR *P. aeruginosa* include using lytic bacteriophage or pyocins, or small molecules that target quorum sensing, biofilm formation, type III secretion, or exotoxin to impair pathogenesis (2). The secreted metalloprotease LasB has stood out among the virulence factors targets, with multiple efforts underway to develop small-molecule inhibitors against it, summarized in (3). It is an attractive target due to its conservation, essentiality, and direct contribution to pathogenesis and pathology through the cleavage of numerous host proteins. These include cleavage and loss of function in tight-junction cadherins, matrix proteins (collagens, elastin, and laminin) and immune effectors (cathelicidin, complement proteins, immunoglobulins, lysozyme, transferrin) and the direct activation of inert IL-1β (3–5). The sum of these targets promotes bacterial replication, invasion, and pathological inflammation, altogether contributing to the morbidity and mortality observed during infection. Therefore, as an adjunctive therapeutic, inhibitors of LasB have the potential to ameliorate these complications and promote effective clearance by immune effectors and protect tissue function.

We recently repurposed the investigational matrix metalloprotease inhibitors Marimastat and Ilomastat as inhibitors of LasB (5), suggesting drug repurposing may have therapeutic utility. To-date, the only metalloprotease inhibitor in common use is Periostat, an oral formulation of low-dose doxycycline sufficient to inhibit the activity of metalloprotease activity of human collagenase (MMP-1), but which does not achieve antimicrobial activity (6, 7). In this study we investigated whether doxycycline and other structurally-related antibiotics of the tetracycline family inhibit the *Pseudomonas* protease, LasB. We show that tetracyclines inhibit LasB activity, and at sub-antimicrobial concentrations, sensitize the bacterium to neutrophil killing. In a murine model of pulmonary infection with an MDR clinical isolate fully resistant to tetracyclines, we find *in vivo* inhibition of LasB by tetracycline and doxycyline that limits pathology. This work highlights new therapeutic strategies for targeting inflammation and proteolysis, and may have utility in the management of complicated disease by targeting metalloprotease virulence factors common among pathogens.

## MATERIALS AND METHODS

### Bacterial strains

*P. aeruginosa* MDR4, a strain resistant to all antibiotic but colistin,has been previously described (5). Bacteria were routinely propagated in Luria broth (LB) (Hardy Diagnostics) or on LB agar plates at 37 ºC. Bacteria were grown to late exponential phase (OD_600_ 1.2) in LB then washed and diluted in PBS for infections.

### LasB purification

pET-His-LasB-CT_(197-301)_ (Genbank: CP129519.1) was created using PIPE cloning with primers: 5’ TCTGTTCCAGGGGCCCGCCC**ATG**GCCGAGGCGGG and 3’ TGCTCGAGTGCGGCCTTACAACGCGCTCGGG. Protein expression in *Escherichia coli* BL21 (DE3) was induced at 18 °C overnight with 0.2 mM isopropyl-β-d-thiogalactopyranoside (IPTG) when OD_600_ reached 0.6. Cells were collected and resuspended in PBS and homogenized by ultrasonication. His-LasB_CT_ was purified by affinity chromatography using HisPur Nickel Resin (Thermo Scientific).

### Molecular docking

The mature LasB sequence of PDB: 6F8B, and the chemical structures of each antibiotic from PubChem using the Chai-1 Model (8). Doxycycline was similarly modeled with active human MMP1 from the crystal structure PDB: 3SHI. Modeling was agnostic with no input restraints specified, and the best scoring model was selected for analysis in ChimeraX. Matchmaker was used to orient each molecule to the same position, displaying antibiotic occupancy in the substrate cleft of the enzyme, and surface electrostatics displayed in conventional red-blue coloring.

### Inhibition of LasB by tetracyclines

A peptide of the sequence HDAPVRSLN internally quenched by labeling with Methoxycourmarin (Mca) and dinitrophenyl (Dnp) on C- and N-terminus respectively, was used as a LasB-reporter as in previous studies (9, 10). 80 nM LasB was incubated in PBS, 0.01% Tween-20, with 2 µM of HDAPVRSLN peptide. Doxycycline (RPI), tetracycline (Alfa Aesar), minocycline (Cayman Chemical), or sancycline (Cayman Chemical) were included at the indicated concentrations to test inhibition, compared to no antibiotic control. 0.1% phenylmethylsulfonyl fluoride (Sigma) was added to evaluate LasB-specific protease activity for *in vitro* and *in vivo* studies to reduce background (5). The reaction was continuously monitored in a black 96-well plate (Costar) using a Victor plate reader (PerkinElmer). Fluorophore excitation was at 323 nm and emission at 398 nm and the maximum kinetic velocity was calculated as previously (9).

### Neutrophil isolation and infection

Blood was collected from healthy male and female human volunteers under informed consent with approval by the Emory University Institutional Review Board. Neutrophils were isolated from heparinized blood using PolyMorphPrep (Axis-Shield) as previously described (11). Cell viability, purity, and concentration were assessed microscopically using 0.04% trypan blue, then suspended in RPMI lacking phenol red, fetal bovine serum, or antibiotics. Except when noted, donor neutrophils and cultured cells were routinely infected by co-incubation with *P. aeruginosa* at a multiplicity of infection of 10, spun into contact for 3 min at 300 g, then infection allowed to proceed 2 h during incubation 37°C in 5% CO_2_.

### Animal Experiments

All animal experiments were performed following Association for Assessment and Accreditation of Laboratory Animal Care (AAALAC) and the Office for Laboratory Animal Welfare (OLAW) guidelines and were approved by the Institutional Animal Care and Use Committees of UCSD or Emory University. Eight-to-ten week old male and female C57Bl/6 mice were anesthetized with ketamine/xylazine intraperitoneally, then 5 x 10^6^ CFU MDR4 inoculated intratracheally in 30 µl of 1x PBS. For antibiotic treatment, 1 µg/g Doxycycline or 1 µg/g Tetracycline were administered intraperitoneally at time of infection. These are approximately half the sub-antimicrobial dose against susceptible organisms established in previous studies (1.75 µg/g) (12). At 24h post-infection mice were euthanized by CO_2_ asphyxiation, and bronchiolar lavage fluid was harvested. White blood cells in the bronchiolar lavage fluid were counted on a hemocytometer, while cytologic examination was performed on Cytospin preparations fixed and stained using Hema 3 (Fisher HealthCare). Histologic sections were prepared from formalin-fixed and paraffin-embedded lungs, stained with hematoxylin and eosin (H&E) and imaged on a Nanozoomer 2.0Ht Slide Scanner (Hamamatsu). LasB protease activity in the lavage was assessed by incubation with the peptide as described above.

### Statistics

Statistical analyses were performed using Prism 10 (GraphPad) using 1-way or 2-way t-test or ANOVA as appropriate. No blinding or exclusion criteria were applied. Tukey post-tests were used to correct for multiple comparisons. Statistical significance is indicated as (*, P < 0.05; **, P < 0.005; ***, P < 0.0005; ****, P < 0.00005). All error bars show the mean and the standard deviation (s.d.). Data are representative of at least three independent experiments.

## RESULTS

Given the possibility of cross-inhibition of metalloproteases across species (5), we sought to examine whether additional known metalloprotease inhibitors could inhibit the Pseudomonal protease virulence factor LasB. First, since doxycycline is known to have broad and nonspecific discrimination across matrix metalloproteases, we used molecular structure prediction modeling to examine the predicted interaction with the major human collagenase MMP1. While clinical efficacy is well-established across numerous studies (13), the molecular basis for inhibition is not established. As typical of protease inhibitors, doxycyline was predicted by unguided analysis to bind within the catalytic pocket (**Fig 1A**) and interact with the histidines required for coordinating the catalytic zinc, a mechanism shared with designed peptidomimetic metalloprotease inhibitors (13). LasB similarly has a substrate binding cleft with surrounding surface electrostatics determining substrate specificity, gating the catalytic site (**Fig 1A**). Doxycycline, as well as tetracycline, minocycline, and sancycline were all predicted to occupy the substrate pocket in a manner that would similarly inhibit catalysis (**Fig 1A**). Therefore, we examined their ability to inhibit LasB cleavage of the internally-quenched LasB substrate HDAPVRSLN, established previously (5). At 60 µg/ml, all four tetracyclines significantly reduced the proteolytic activity of purified LasB relative to negative control (**Fig. 1B**).

**Figure 1.**
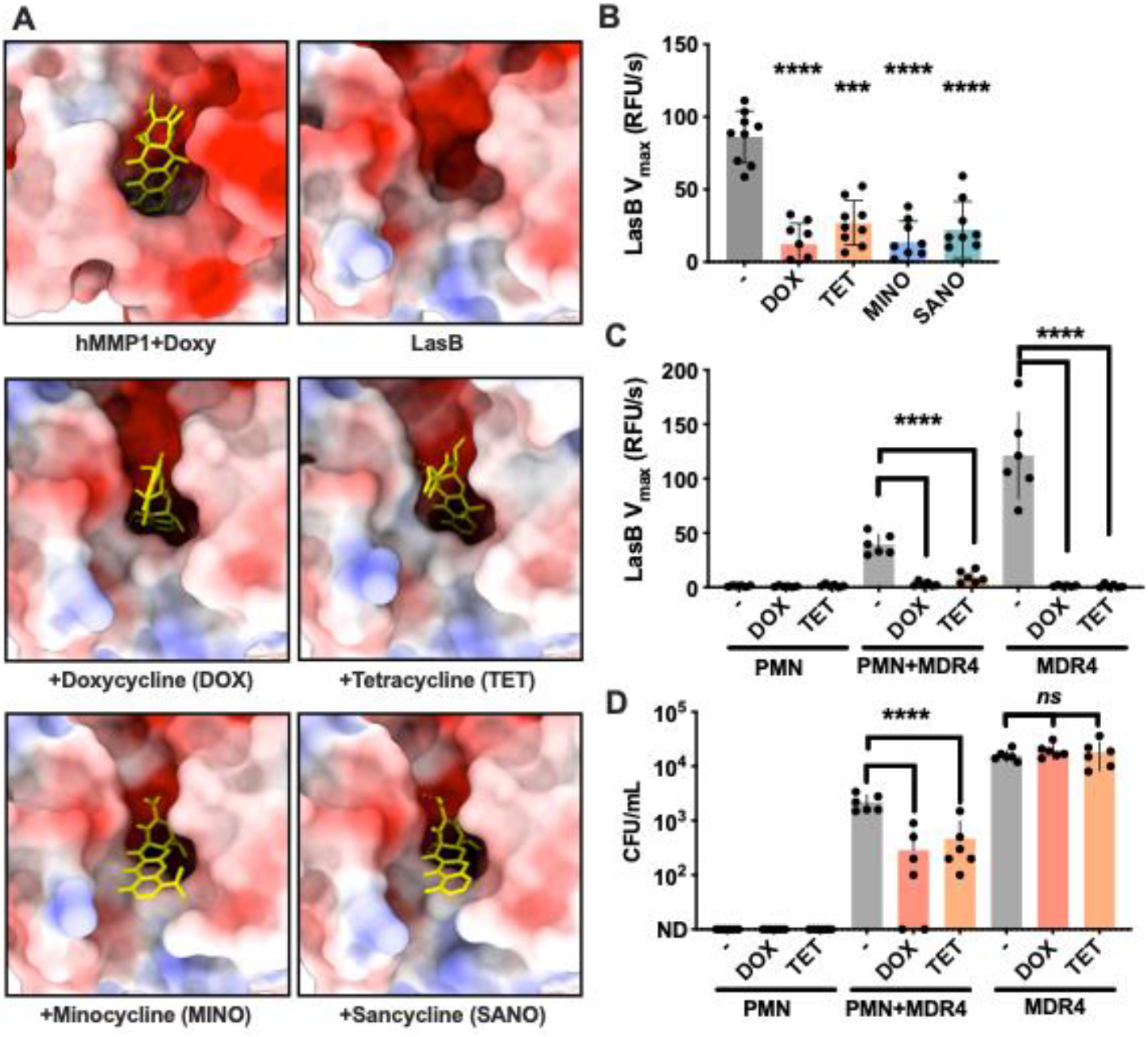
Inhibition of LasB by tetracyclines. (**A**) Models of the substrate pocket of human MMP1 or *Pseudomonas* LasB, with the indicated antibiotic (yellow). Protein surface electrostatics are colored red and blue for negative and positive charges, respectively, and white color represents neutral residues. (**B**) Cleavage of HDAPVRSLN peptide after 30 min incubation with purified LasB in the presence or absence of tetracycline-family antibiotic indicated. (**C**) Cleavage of HDAPVRSLN peptide after 2 hr infection of neutrophils by *Pseudomonas* MDR4, or neutrophils or bacteria alone, in the presence of the indicated antibiotic or control. (**D**) Colony count of *Pseudomonas* MDR4 in the infection of (C). *\*\*\*P<0*.*0005*; *\*\*\*\*P<0*.*00005* (ANOVA; Tukey post-test). Experiments were performed three times; error bars represent standard deviations.

Numerous studies establish the importance of neutrophils in controlling *P. aeruginosa* infection, and contributions of LasB to killing (5). We modeled this interaction *in vitro* with neutrophils collected from healthy human donors and infected with *P. aeruginosa* MDR4, a multidrug resistant clinical isolate resistant to all classes of antibiotics except colistin (5). Neutrophils alone had no activity against the LasB substrate, and this was unimpacted by the presence of doxycycline or tetracycline (**Fig. 1C**). Cultures of MDR4 in the same media rapidly cleaved the LasB substrate, and this was fully inhibited by doxycycline or tetracycline (**Fig. 1C**). When the bacteria were incubated with neutrophils, there was a decline in LasB proteolysis relative to media alone, but which was still fully inhibited by both doxycycline and tetracycline (**Fig. 1C**). When the surviving bacteria were plated in this experiment, neither doxycycline nor tetracycline impacted the survival of MDR4 in culture (**Fig. 1D**). However, both antibiotics decreased bacterial survival during neutrophil infection (**Fig. 1D**). Additionally, even in the absence of antibiotics, neutrophils reduced bacterial titer relative to media-only control (**Fig. 1D**), providing an explanation for why less LasB was measured in this condition (**Fig. 1C**). Together these data suggest that tetracyclines can promote killing of *P. aeruginosa* by immune cells, even against antibiotic-resistant strains.

To validate our proof-of-concept for inhibiting LasB-mediated inflammatory pathology with antibiotics, we examined inflammation in mice infected with MDR4 as previously (5). In a pulmonary infection model, MDR4 induced significant pathology (**Fig. 2A**), with marked immune cell infiltration into the airway as seen in human disease (**Fig. 2B**). This was reduced by treatment with doxycycline or tetracycline (**Fig. 2A, 2B**). By lavage, significant reductions in active LasB were detected in the infected lungs of mice treated with doxycycline or tetracycline (**Fig. 2C**). This reduced inflammation and neutrophil recruitment is suggestive of a protective effect beyond antimicrobial activity that ultimately contribute to preserving the pulmonary architecture.

**Figure 2.**
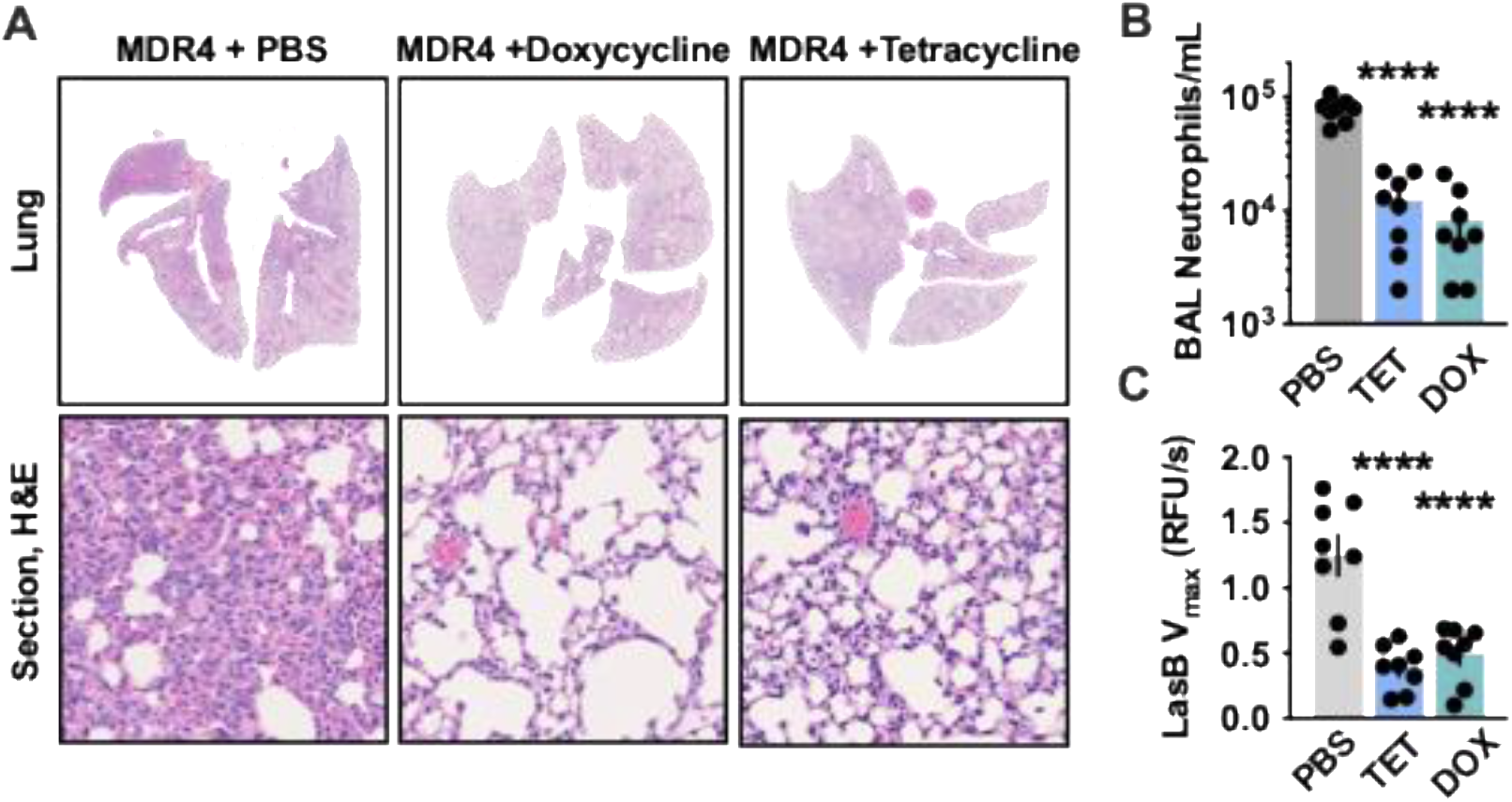
Tetracyclines prevent pathological inflammation during *Pseudomonas* lung infection. C57BL/6 mice were intratracheally infected with *Pseudomonas* MDR4 and treated with Doxycycline, Tetracycline, or PBS control, and the infection was allowed to proceed 24 h until euthanasia. (**A**) Representative histology sections cytological smears of bronchoalveolar lavage fluid prepared with differential MGG stain. Mouse bronchiolar lavage was harvested and (**B**) neutrophils enumerated, or (**C**) cleavage of HDAPVRSLN peptide measured over 30 min incubation. Data are pooled from two independent experiments, N=8, error bars represent standard deviations. *\*\*\*P<0*.*0005*; *\*\*\*\*P<0*.*00005* (ANOVA; Tukey post-test).

## DISCUSSION

*P. aeruginosa* destroys the lung architecture and function through microbe-, inflammation-, and neutrophil-mediated degradation of mucin layers and structural proteins of the pulmonary connective tissue. Uncontrolled proteolysis directly correlates with disease severity (14), confounds effective antimicrobial responses, and may ultimately contribute to the decline in lung function. Additional important considerations revealed by this works are the effect of antibiotics beyond that on the microbe itself, but on metalloprotease virulence factors, as well as on host metalloproteases, and host signaling pathways (12).

Notably this was achieved at concentrations below those that are antimicrobial, and lower than those that inhibit other proinflammatory processes (12).

LasB is typically expressed during acute infections and in murine lung infections model and is essential in these conditions (10). Some clones isolated from patients chronically infected by *P. aeruginosa*, particularly from persons where effective clearance is impaired by cystic fibrosis, have mutations preventing high level LasB expression (3). The prevalence of infections by populations that are fully LasB non-expressors is likely low, but through the cross-inhibition of matrix metalloproteases and other processes contributing to pathological inflammation and antiinfective failure, so some therapeutic effect may yet remain from tetracyclines. Additionally, differential bioavailability at each potential infection site can impact the therapeutic utility of LasB inhibition. For example, standard oral treatments can reach over 150 ug/ml (doxycycline) and 300 ug/ml (tetracycline) in the urine, suggesting the possibility this strategy could be useful in the treatment of non-systemic *P. aeruginosa* UTIs (15). Lastly, while tetracyclines are recognized as generally safe in adults, mild side effects can be evident with longer-term use. This would need to be a consideration for its incorporation as antibiotic or adjunctive therapeutic.

## Acknowledgments

We thank the members of LaRock lab for helpful discussions. We appreciate the technical support provided by the Children’s Healthcare of Atlanta and Emory University’s Children’s Clinical and Translational Discovery Core for whole blood and cell processing. Molecular graphics and analyses performed with UCSF ChimeraX, developed by the Resource for Biocomputing, Visualization, and Informatics at the University of California, San Francisco, with support from NIH R01-GM129325 and the Office of Cyber Infrastructure and Computational Biology, NIAID. C.N.L. is supported by National Institutes of Health grants R01-AI153071 and R01-AI180089. C.N.L. is a Burroughs Wellcome Fund Investigator in the Pathogenesis of Infectious Disease. S.H. received support from the SOAR program of the Laney Graduate School, Emory University. The content of this publication is solely the responsibility of the authors and does not necessarily represent the official views of any of its funders. No funders contributed to the study design or conclusions.

## Author contributions

M.C., J.S., S.C., D.L., J.B., and C.L., designed experiments and interpreted the data. S.C., D.L., J.S., and C.L. conducted the studies. M.C. and C.L. wrote the manuscript with the assistance of all the authors. All authors approved the final manuscript.

